# Efficient direct and limited environmental transmission of SARS-CoV-2 lineage B.1.22 in domestic cats

**DOI:** 10.1101/2022.06.17.496600

**Authors:** Nora M. Gerhards, Jose L. Gonzales, Sandra Vreman, Lars Ravesloot, Judith M.A. van den Brand, Harmen P. Doekes, Herman F. Egberink, Arjan Stegeman, Nadia Oreshkova, Wim H.M. van der Poel, Mart C.M. de Jong

## Abstract

Susceptibility of domestic cats for infection with SARS-CoV-2 has been demonstrated by several experimental studies and field observations. We performed an extensive study to further characterize transmission of SARS-CoV-2 between cats, both by direct contact as well as by indirect contact. To that end, we estimated the transmission rate parameter and the decay parameter for infectivity in the environment. Using four groups of pair-transmission experiment, all donor (inoculated) cats became infected, shed virus and seroconverted, while three out of four direct contact cats got infected, shed virus and two of those seroconverted. One out of eight cats exposed to a SARS-CoV-2-contaminated environment became infected but did not seroconvert. Statistical analysis of the transmission data gives a reproduction number *R_0_* of 2.18 (95% CI: (0.92-4.08), a transmission rate parameter β of 0.23 day^-1^ (95% CI: 0.06-0.54), and a virus decay rate parameter μ of 2.73 day^-1^ (95% CI: 0.77-15.82). These data indicate that transmission between cats can be sustained (*R_0_*>1), however, infectiousness of a contaminated environment decays rapidly (mean duration of infectiousness 1/2.73 days). Infections of cats via exposure to a SARS-CoV-2-contaminated environment cannot be excluded if cats are exposed shortly after contamination.

## Introduction

Since the first cases of atypical pneumonia due to SARS-CoV-2 were identified in December 2019, a large number of animal studies were performed to assess which species are susceptible to SARS-CoV-2 infection. Next to susceptibility to SARS-CoV-2, it is relevant to determine which species develop clinical disease and which species can transmit the virus. In several studies, animals were experimentally inoculated with high doses of SARS-CoV-2 to demonstrate species susceptibility, such as ferrets, cats, hamsters, non-human primates, rabbits and other species (reviewed by Jo et al. (2021), for instance). Next to these experimental infections, a number of animals were shown to be infected naturally by contact with infected humans, such as minks (Oreshkova et al. 2020), dogs (Barroso-Arevalo et al. 2021), and both domestic and big cats (Patterson et al. 2020; Rotstein et al. 2022).

Associated with species susceptibility to infection and/or disease, it is relevant to assess the risk of SARS-CoV-2 transmission between animals and humans. This is of particular concern for animal species that live in close contact with humans. These animals can then be a reservoir and/or an intermediate host for humans. There are a few experimental studies addressing transmission between cats, showing that infected cats can pass on the virus by aerosol or contact transmission to other cats (Halfmann et al. 2020; Gaudreault et al. 2020; Shi et al. 2020; Bao et al. 2021). A commonality of these studies is an inoculation dose above 10^5^ PFU/TCID_50_, housing of animals in smaller isolators/cages and frequent sampling under anaesthesia. A systematic review of these experimental data combined with household data confirmed that sustained direct contact transmission between cats is probable (R_0_> 1) and that the risk of indirect transmission via aerosols is possible, although less likely than direct contact transmission (Gonzales et al. 2021).

The risk of transmission via exposure to contaminated surfaces (environment) has not yet been studied. Assessing this risk is relevant because susceptible cats could be exposed to contaminated surfaces during visits at (holiday) pet shelters or veterinary clinics if proper hygienic measures are not implemented. In addition, successful infection of cats exposed to an environment contaminated with SARS-CoV-2 by other cats could be an indicator of the potential risk for cat-to-human transmission via a contaminated environment. Whilst transmission of SARS-CoV-2 from humans to animals has been well-documented, reports of transmission from animals to humans remain limited. To date, mink-to-human transmission has been confirmed (Oude Munnink et al. 2021) and recent events have highlighted potential transmission from infected hamsters to humans in a pet-shop in Hong Kong (Yen et al. 2022). Moreover, a recent report describing a possible cat-to-human transmission of SARS-CoV-2 highlights the possibility of this transmission pathway (Sila et al. 2022), which is a relevant pathway given the proximity between humans and pets.

To assess the potential risk of cats for spreading SARS-CoV-2 via contaminated environment and provide further evidence of direct contact transmission we have performed a transmission experiment in domestic cats after infection with SARS-CoV-2. We confirm that direct contact transmission of SARS-CoV-2 is efficient between cats and that the infectiousness of a contaminated environment decays rapidly, limiting thereby the efficiency of environmental transmission alone. Nevertheless, the indirect, environmental transmission route may still be possible.

## Results

The study design is explained in detail in the Methods section and summarised in Figure 1. Four independently housed pairs of cats were used to assess direct transmission and contamination of the environment (pen) where new naive cats (two per contaminated pen) were introduced following removal of the pair of cats used to assess direct transmission. The sample size was calculated based on data retrieved from literature (Gonzales et al. 2021). From the direct transmission pairs, the four donor cats became infected following inoculation and three of those transmitted infection to their corresponding contact cats. Of the eight cats exposed to the contaminated pens, only one got infected. Below we provide detailed results of the observed infection characteristics and the quantitative assessment of transmission.

**Figure 1.**
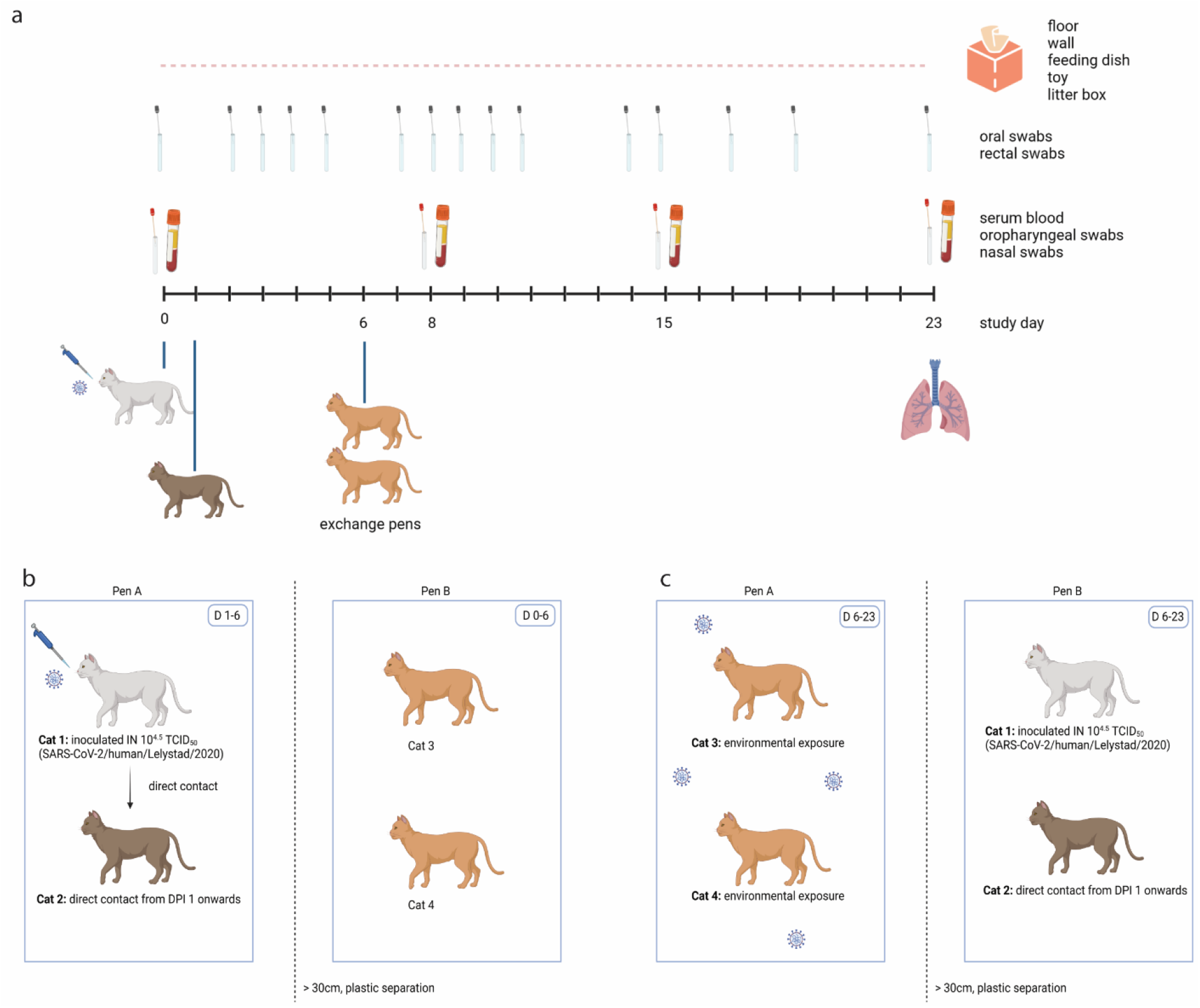
Experimental design of one out of four replicate groups. **(a)** Schematic timeline. One cat was inoculated intranasally on D0, and brought in contact with a naïve cat on D1 in pen A. On D6, these two cats were exchanged with two other naïve cats. Blood samples, oropharyngeal swabs and nasal swabs were collected under general anaesthesia on D0, D8, D15 and upon euthanasia. Oral and rectal swabs were collected more frequently without anaesthesia. Environmental samples were collected daily, and all cats were euthanized on D23 except of one cat who died before the end of the study by a cause not related to SARS-CoV-2. **(b)** Until D6, cat 1 (inoculated donor cat) and cat 2 (direct contact recipient cat) were housed in pen A and contaminated the environment. Cat 3 and cat 4 were housed in pen B, which was separated from pen A by a plastic separator and space. **(c)** From D6 onwards, cats 3 and 4 were housed in contaminated pen A. Cats 1 and 2 were housed in pen B.

### Clinical signs and pathology

Two of the 4 inoculated cats occasionally showed mild serous nasal discharge. All 12 cats having direct contact or indirect contact with the inoculated cat remained without clinical signs. The body weights of all 16 cats remained constant. A video-based analysis of activity of the direct transmission pairs revealed no changes in activity post inoculation compared to before inoculation (baseline measurement), indicating that inoculated and direct contact cats displayed a similar activity pattern regardless of SARS-CoV-2 inoculation/direct exposure (Supplemental material 1). One cat of group 4 (cat 4.3) died on D18 and was necropsied the same day. It was confirmed that the death was unrelated to SARS-CoV-2 infection. All other cats were euthanized on D23. Upon necropsy, no lesions were observed in gross pathology and there were no substantial differences in relative lung weights between the animals (data not shown). Lung tissue showed no SARS-CoV-2-related histopathology, however, mild lung changes such as lymphoplasmacytic bronchoadenitis, bronchus associated lymphoid tissue (BALT) hyperplasia, infiltrates of macrophages and few neutrophils in alveolar lumina were observed in inoculated animals, but also in direct and indirect contact animals (data not shown) suggested to be nonspecific. Other evaluated organs (nasal conchae, trachea, tracheobronchial lymph node, duodenum, ileum, colon, mesenterial lymph node and pancreas) showed also no substantial histopathological findings. No viral antigen could be detected by immunohistochemistry in any of the investigated tissues (data not shown).

### Viral RNA load in swabs and organs

Oral and rectal swabs were taken frequently, while nasal and oropharyngeal swabs were sampled only on a few time points (D0, D8, D15 and D23). Swabs were analysed by both total E-gene PCR according to Corman et al. (2020), and by subgenomic PCR (sgPCR) according to Wolfel et al. (2020).

SARS-CoV-2 RNA was detected by both total E-gene PCR as well as sgPCR in multiple swabs of all inoculated cats (X.1) from D2 until the end of the study (Figure 2a). In swabs of all four direct contact cats (X.2), SARS-CoV-2 total E-gene RNA was detected, while only three cats (1.2, 2.2 and 4.2) also had at least one positive sample in the sgPCR. Of the indirect contact cats (X.3 and X.4) exposed to contaminated environment, total E-gene RNA was detected in swabs in groups 1, 3 and 4, while in group 2, both cats remained negative for SARS-CoV-2 RNA (Figure 2b). Of all swabs collected from all indirect contact cats, only one sample tested positive in the sgPCR (oral swab of cat 1.3 on D7).

**Figure 2.**
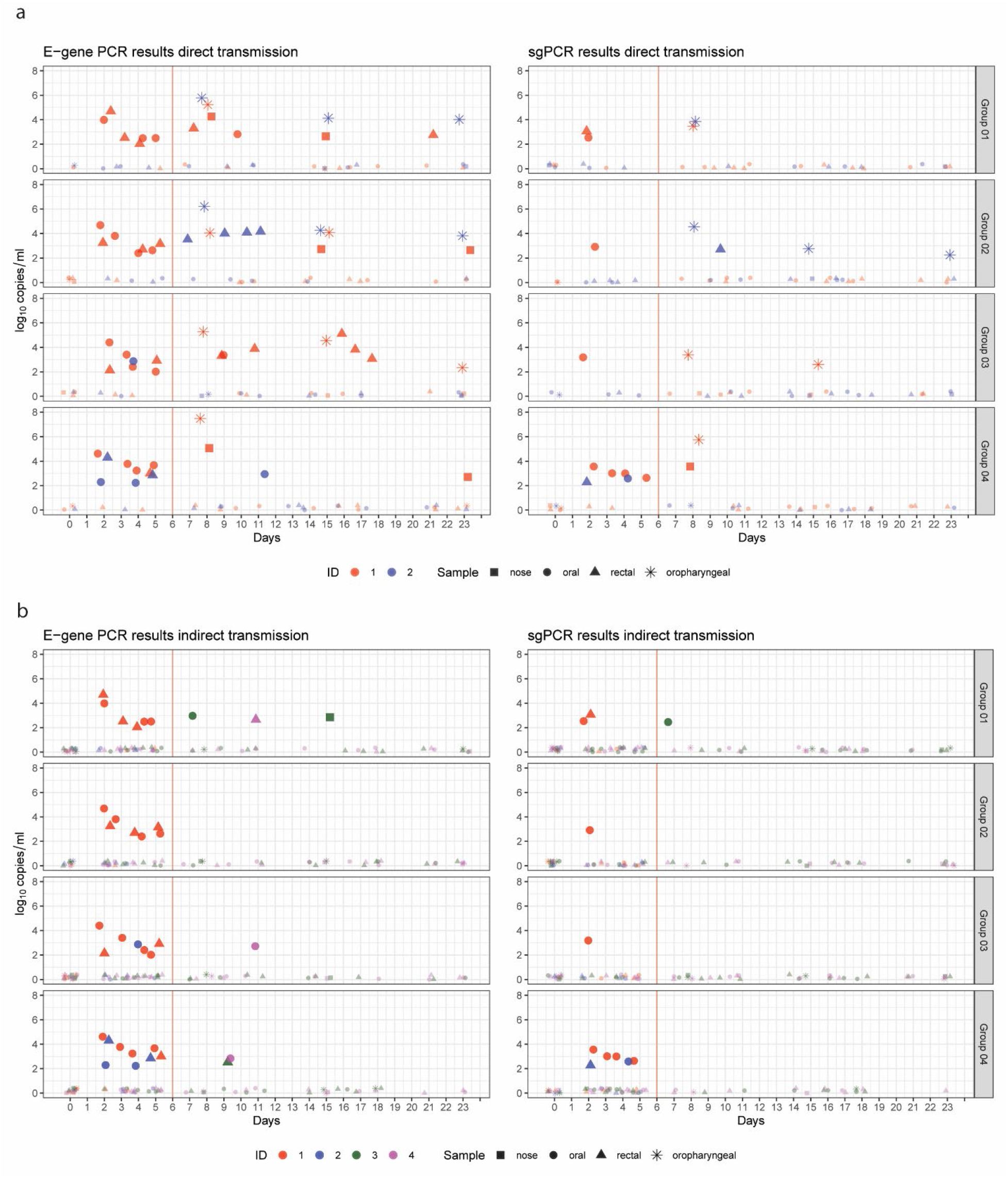
E-gene and subgenomic PCR results on swabs. For total E-gene PCR, the mean log10 RNA copies/mL from two technical replicates of the same swab are shown. The replicates were generated by isolating RNA from the swab sample in duplicate and subsequent PCR. The sgPCR was performed once with the RNA sample from the replicate that showed a lower Ct value in the total E-gene PCR. Shapes indicating the results are jittered so that overlapping shapes can still be observed. The red line indicates the day of removal of cats 1 and 2 from pen A and housing them in pen B. **(a)** Swabs from the direct transmission pairs. **(b)** Samples on the left of the red vertical line show results from direct-transmission pairs, cats 1 and 2 [this is a repetition (jittered) of the data shown for days 0 to 6 in panel a)], while the samples on the right of the red vertical line are collected from cats 3 and 4 (indirect transmission cats) in pen A.

After necropsy, respiratory organs (nasal conchae, trachea and lungs) of all cats were analysed by total E-gene PCR. All samples were PCR-negative except of nasal conchae of cats 1.1, 2.1 and 4.1 (3 of the 4 inoculated cats). Noteworthy, these samples were collected 23 days post inoculation.

### Serology and immunology

Sera and heparinized blood samples were collected on D0, 8, 15 and 23 of all cats. Virus neutralizing antibodies were detected in all inoculated cats (X.1) on D8, 15 and 23 with neutralizing titres ranging from 90 to 810 (Figure 3a). Also, anti-S1 antibodies were detected in all inoculated cats by ELISA (Figure 3b). Two out of the four direct contact cats (1.2 and 2.2) developed neutralizing (titre range 5.7 to 155) and anti-S1 antibodies, while the other two direct contact cats (3.2 and 4.2) did not seroconvert. None of the indirect contact cats developed a serological response.

**Figure 3.**
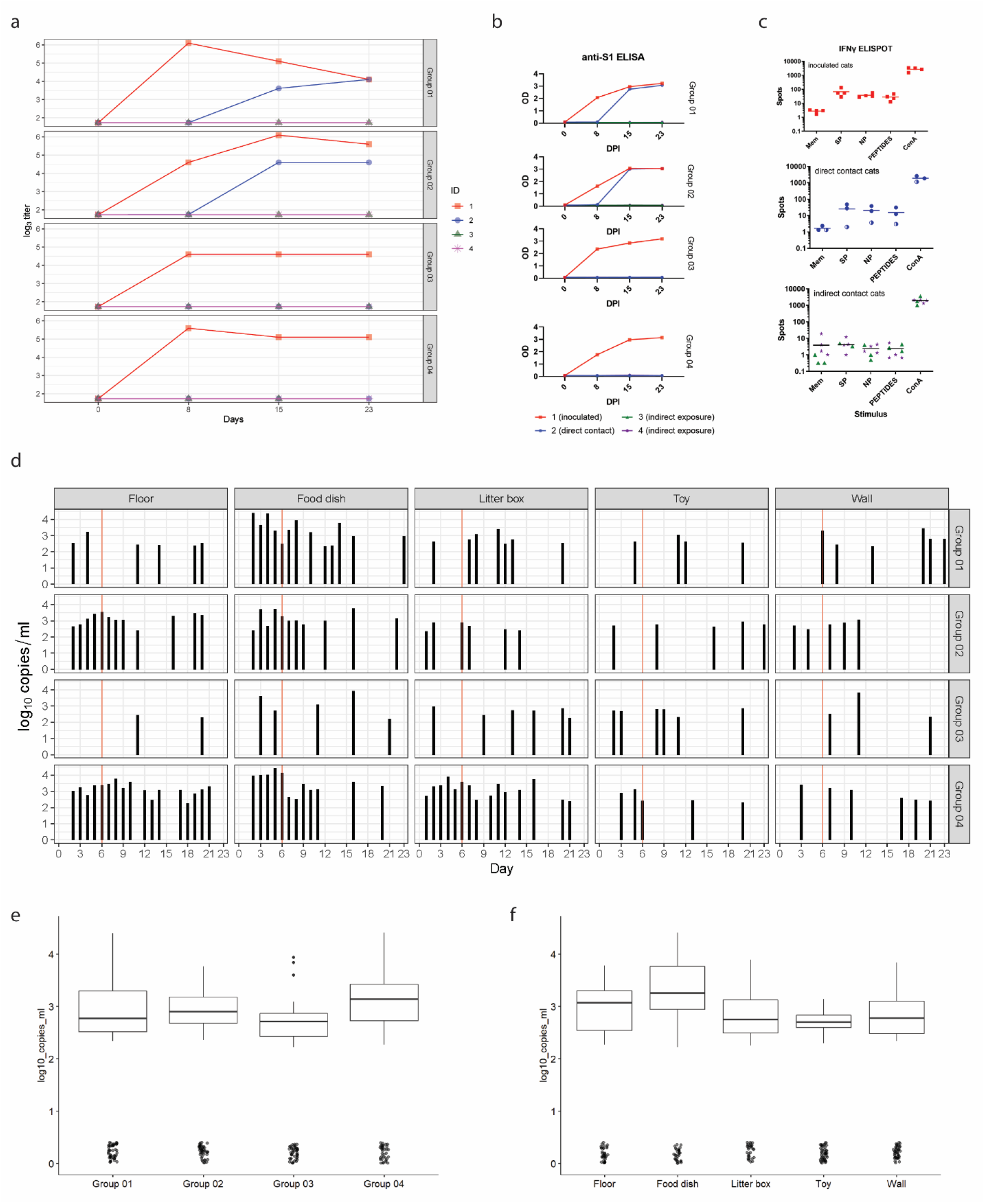
Serology, immunology and environmental contamination. **(a)** Neutralizing antibody titres expressed as 50% neutralization titre (MN50). All four inoculated cats (cats X.1) and two of the direct contact cats (cats 1.2 and 2.2) developed neutralizing antibodies, while the indirect exposure cats (cats X.3 and X.4) did not seroconvert. **(b)** Anti-S1 antibody response measured by ELISA at OD420. Antibodies directed against S1 were detectable in all inoculated cats, as well as the direct contact cats in group 1 and 2. **(c)** Interferon-γ response of PBMCs after stimulation for 48h with spike protein (SP), nucleoprotein (NP), a peptide pool (PEPTIDES), and Concanavalin A (ConA) and medium (Mem) as control, measured by ELISpot analysis. An interferon-γ response was observed in all inoculated cats and direct contact cats in group 1 and 2 after stimulation with SP, NP, the peptide pool and ConA. No interferon-γ response was detected in the direct contact cat of group 4 (indicated by half-filled bullets) or in the indirect contact cats, except of for the positive control ConA. Cats 3.2 and 4.3 were not tested due to an insufficient number of PBMCs. **(d - f)** Environmental samples analysed by E-gene PCR. **(d)** Overall viral RNA levels of the environment per group and sample type. Viral RNA was detectable throughout the entire study period. **(e)** Distribution of viral RNA load in the different groups. **(f)** Distribution of viral RNA load per sample type.

To evaluate memory T-cell interferon-γ production, peripheral blood mononuclear cells (PBMCs) were stimulated for 48h with a SARS-CoV-2 spike (S), nucleoprotein (NP), or a SARS-CoV-2 spike protein peptide pool, and analysed with an ELISpot assay (Figure 3c). Compared to medium stimulation, a clear interferon-γ response (increased number of spots) was measured in all four inoculated cats and two direct contact cats (1.2 and 2.2). This indicates that these cats were able to mount a specific cellular immune response on DPI 23 (inoculated cats) or between DPI 17-22 (direct contact cats). In none of the indirect exposure cats, a memory T-cell interferon-γ response was observed (X.3 and X.4). For cat 3.2 and 4.3 we obtained insufficient PBMCs to perform an ELISpot assay.

### Environmental contamination

Environmental samples (floor, wall, litter box, feeding tray and toy) were collected daily from all pens A, and analysed by total E-gene PCR. Viral RNA could be detected in all types of samples until the end of the study (Figure 3d). There was variation between the groups, amongst which the highest median genome copy numbers in the different sampled surfaces were found in group 4, and of all sample types, the feeding dishes were those samples with the highest median genome copy numbers (Figure 3e and 3f). The average temperature and relative humidity (RH) in the experimental pens during the course of the experiment were 19.65°C and 54.85%, respectively.

### Statistical analysis of transmission

For transmission to be considered successful we defined contact cats as being infected and infectious either based on the sgPCR or on seroconversion. For quantification of the transmission parameters, the moment the cats were identified as infected (moment of infection) and infectious was based on the E-gene PCR (evidence of exposure) because of the higher sensitivity of this PCR; i.e. the moment of infection was considered to be 1-2 days before the first E-gene-positive sample (latent period). They were then counted as infectious from the first day until the last day that they were observed as E-gene-PCR positive in any of the samples tested.

In Table 1 the maximum likelihood estimates of the parameters describing the transmission process are given for four different analyses using an extension of the direct transmission SIR model with environmental exposure. The analysis is done on four different scenarios which are represented as datasets. Datasets for scenarios ‘data1’ and ‘data2’ are based on using seroconversion as primary indicator for being infected and infectious, while ‘data3’ and ‘data4’ make use of the sgPCR results as primary indicator. ‘Data1’ and ‘data3’ uses only data from the direct transmission pairs, while ‘data2’ and ‘data4’ also included the data from the environmental exposure pairs used to assess indirect transmission via the environment (see supplementary material 3 for all four datasets). The results of the analysis using data1 illustrates why the environmental SIR model is used. In this experiment, the direct-contacts become infected very late in the experiment and also two cats escape infection, indicating that build-up of infectiousness was required for infection to occur. This does not fit the direct transmission SIR model which assumes that direct transmission infection would occur with equal probability on each day of the exposure. Hence the environmental SIR model fits the data only with a low decay rate parameter. This estimate implies that, after the direct exposure period ends, there would still be high infectivity and cats added to the contaminated environment would readily get infected. However, when we add the indirect exposure period to the data we see that none of the eight indirect exposed cats seroconverted. This is in spite of the presence of E-gene-PCR-positive material in the environment and in samples from the exposed cats for the whole experimental period (two extra weeks) (Figure 2 and 3). Using this dataset (data2) the best fitting model estimates the decay rate parameter as very large, making the environmental SIR equivalent to the direct transmission SIR model.

**Table 1.** Results for the stochastic SIR model with environmental transmission. Four different scenarios (datasets) were analysed based on the observed infection events using either seroconversion (‘sero’) or sgPCR (‘SG’) as test to determine which cats had become infected and analysing only the first part of the experiment with direct exposure (‘direct’) or also the data with indirect exposure (‘both’). The model yielded two parameters: the transmission rate parameter β and the decay rate parameter μ. From those parameters the reproduction ratio of SARS-Cov-2 in cats was calculated [in this case based on an infectious period of 6 days according to Gonzales et al. (2021)].

| Scenario | Test | Data period | $\beta$ (day<sup>-1</sup>) | $\mu$ (day<sup>-1</sup>) | R |
| --- | --- | --- | --- | --- | --- |
| data1 | sero | direct | 0.044<br>(0.013-0.105) | 0.001 | 528.2<br>(156.1-1260.4) |
| data2 | sero | both | 0.21<br>(0.06-0.49) | ∞ | 1.26<br>(0.37-3.02) |
| data3 | SG | direct | 0.41<br>(0.16-0.85) | ∞ | 2.50<br>(0.97-5.15) |
| data4 | SG | both | 0.23<br>(0.06-0.54) | 2.73<br>(0.77-15.82) | 2.12<br>(0.92-4.08) |

For the analysis performed using the scenario represented in the datasets ‘data3’ we observed that the direct transmission SIR model fits the data best because the direct-contact cat 4.2 (Figure 2) became infected very early during exposure at the interval 1 to 2 days. However, when using the sgPCR as indicator of infection, there is an infection observed (cat 1.3) in the indirect transmission group (‘data4’). Using this scenario (data4), we get a finite estimate of the decay rate parameter. The maximum likelihood estimate (Table 1) is calculated for the transmission rate parameter β = 0.23 day^-1^ (95% confidence Intervals CI: 0.06-0.54) and for decay rate parameter μ = 2.73 day^-1^ (0.77-15.82), which implies an average survival time (duration of infectiousness in the environment = 1/ μ) of 8.8 (1.5 – 31.2) hours. The interpretation of these estimates can be seen in Figure 4. For the direct transmission SIR model, infection only occurs during direct exposure, i.e. in the five days the direct contact animals are present. In the environmental SIR model the infection probability increases over time during the direct exposure period and it is still present when the infected animals, contaminating the environment, are removed and naive animals are placed to indirectly expose them to the contaminated environment.

**Figure 4.**
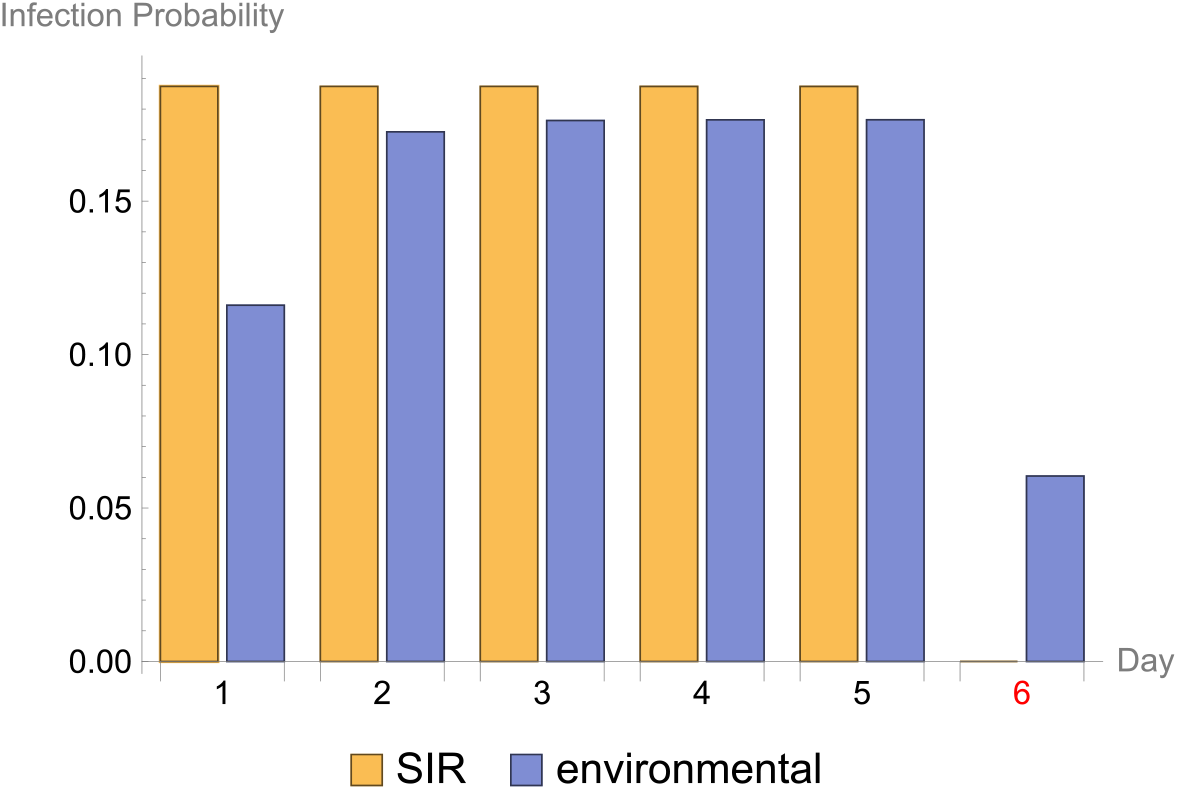
Infection probability. The infection probability for each observed interval in the experiment where the day of the start of the interval is given on the x-axis. The results for two fitted stochastic models are given: ‘SIR’ is the direct transmission SIR model and ‘environmental’ is the environmental transmission SIR model. Day 6 is highlighted in red, because this is the day when the inoculated cat was removed from pen A.

The estimated R_0_ is 2.12 (0.92-4.08) for transmission between cats for the direct and the indirect transmission route combined. This estimate is based on an infectious period of 6 days, which was derived from data available from other transmission experiments in cats (Gonzales et al. 2021).

## Discussion

This study explored direct and indirect transmission of SARS-CoV-2, lineage B.1.22, among domestic cats in an experimental setting. We particularly quantified the duration of infectiousness of an environment with contaminated surfaces. We found that the infectiousness of contaminated surfaces would decay within 8.8 (95%CI: 1.5 - 31.2) hours, making transmission via contaminated surfaces alone inefficient, yet not to be excluded. In addition, we provide further confirmation that cat-to-cat transmission can be sustained.

All four inoculated cats became infected with SARS-CoV-2 both when measured by seroconversion and by sgPCR and remained clinically asymptomatic, except for mild nasal discharge in two cats. Compared to other studies focussing on transmission between cats, we used a lower inoculation dose [0.5-1.5 log lower than Gaudreault et al. (2020); Shi et al. (2020); Bao et al. (2021) or Bosco-Lauth et al. (2020)], and only the intranasal inoculation route. We did this because we hypothesised that a lower dose and intranasal exposure only would be a better approximation to natural infection of the donor cats. This hypothesis is based on published peak viral shedding titers of cats infected by direct contact, which is around 10^4.0^ PFU/mL (Gonzales et al. 2021). The absence of obvious clinical signs, with virus shedding measured in oral, nasal, oropharyngeal and rectal swabs, mild nonspecific histopathological findings, seroconversion of inoculated cats, as well as the transmission to naïve cats by direct contact is in agreement with those other reports. The four inoculated cats transmitted SARS-CoV-2 to two (seroconversion) or three (sgPCR) direct contact cats, and all four direct contact cats had E-gene positive PCR samples.

A major challenge in the analysis of transmission experiments is the definition of ‘being infected’. Almost all cats (except indirect contacts in room 3) were tested positive by E-gene PCR, indicating the presence of viral RNA in samples. These positive samples were further tested by sgPCR, which specifically quantifies the mRNA of the E-gene, generated during virus active replication. Next to the presence of viral RNA or mRNA, the third potential definition of ‘being infected’ is seroconversion, which may be dependent on inoculation dose and follow-up time.

Given the different results in the number of contact cats considered infected based on serology or sgPCR, we analysed transmission using either sgPCR or seroconversion as determinant of successful infection. We explored the use of information generated from only the direct-contact experiments or both direct and indirect contact experiments. Serology as indicator for successful transmission did not provide sufficient information for reliable quantification of the decay rate. Contrasting R_0_ estimates were obtained when using data from direct-contact only (data1) and from direct and indirect contact (data2). However, when using sgPCR as indicator for successful transmission, similar R_0_ were estimated for datasets 3 and 4, which were 2.5 (95%CI: 0.97-5.15) and 2.12 (95%CI: 0.92-4.08), respectively (Table 1). These estimates are in agreement with those made by Gonzales et al. (2021) using published data from direct transmission experiments R_0_ = 3.0 (95% CI: 1.5 – 5.8) or from household infections R_0_ = 2.3 (95%CI: 1.1 – 4.9) and provide further certainty that cat-to-cat transmission can be sustained.

Considering that not all direct contact cats seroconverted even though they tested positive by sgPCR and hence are considered truly infected, our data suggests that surveys focussing on only seroprevalence in cats may be an underestimate of the true prevalence of SARS-CoV-2 in cats. Moreover, discrepancies between samples testing positive on one and negative on another assay can also be explained by varying sensitivities of the assays (Corman et al. 2020; Wolfel et al. 2020).

Although our study contributes to an improved understanding of SARS-CoV-2 transmission amongst cats by providing a detailed estimation of both the transmission rate parameter and the decay rate parameter that both influence transmission, at the same time our study faces some limitations. For instance, we did not follow a potential transmission chain from an inoculated cat to a direct contact and yet another direct contact cat, which would be a sensitive measure for infectiousness of the infected direct contact cats. Next, our follow-up time was limited to 17 days for the environmentally exposed cats, which may have been too short to develop antibodies when cats are exposed to lower amounts of virus compared to an inoculated cat. Furthermore, a sparse sampling under anaesthesia was performed on purpose, however, samples collected from the nose and oropharynx were the most sensitive samples and we may have missed viral shedding of cats. Anaesthesia can negatively impact the immune system (Kurosawa and Kato 2008; Schneemilch, Schilling, and Bank 2004), implying that frequently anesthetized animals may become more sensitive to infections compared to animals infected under natural conditions. Finally, in our analysis we assume that direct contact cats are equally infectious as the inoculated cats. This assumption cannot be tested based on virus excretion because the samples that we took more frequently without anaesthesia (oral swabs and rectal swabs) were less sensitive than those samples taken sparsely under anaesthesia (nasal swabs and oropharyngeal swabs), which would have been more adequate to compare the shedding levels of inoculated and direct contact cats. It also cannot be tested based on transmission to contact cats as only one possibly infected cat (the indirect contact cat) was housed together with a recipient. This did not lead to infection of that recipient cat but no infection also happened in the contact between one out of four inoculated cats and their recipients.

Our analysis allowed the estimation of the expected duration of the infectiousness of a contaminated environment, with the probability of indirect transmission decaying on average within the first 8.8 hours following contamination. This observation contrasts with the levels of RNA found in the assessed contaminated surfaces. Similar levels were observed during the whole experimental period (no apparent decay observed), which confirms that the presence of RNA does not reflect the risk of infection by exposure to contamination. These observations agree with laboratory experiments which showed very low to no environmental decay of RNA for up to 21 days on different surfaces (Paton et al. 2021), while live virus could only be isolated from hours to few days depending on the surface and environmental conditions. i.e. estimated half-lives at room temperature and approximately 40% RH in contaminated stainless steel (same surface as the food dishes) ranged from 6 to 43 hours (Biryukov et al. 2020; Riddell et al. 2020; van Doremalen et al. 2020). While these virus survival studies suggest a potential risk of indirect transmission due to the presence of live virus on contaminated surfaces, our analysis directly assessed the probability of transmission and the duration of this risk. It should be noted that the estimated transmission parameters and duration of infectiousness of a contaminated environment may differ for other variants of SARS-CoV-2, e.g. the omicron variant has been shown to survive longer than other SARS-CoV-2 variants (Hirose et al. 2022), and as well as people, infected animals may shed higher level of virus and the transmissibility of this variant may be higher than the virus variant used for the experiment described here (Zhang et al. 2022).

The estimated decay rate parameter for infectivity is an important finding as it is in line with *in vitro* measurements but not with the decay rate parameter for E-gene positivity in the environment. This finding may help to further study cleaning and biosecurity measures also in humans [e.g. Duives et al. (2021)].

In conclusion, our study provides further evidence for efficient transmission of SARS-CoV-2 between cats by direct contact even when a lower inoculation dose was chosen. In contrast, indirect transmission of SARS-CoV-2 to other cats via contaminated environment is less efficient, although it cannot be excluded.

## Materials and methods

### Ethical statement

The study (experiment number 2020.D-0007.011) was performed under legislation of the Dutch Central Authority for Scientific procedures on Animals (CCD license no. AVD4010020209446), and approved by the Animal Welfare Body of Wageningen University and Research prior to the start of the in-life phase. Animal and laboratory work was performed in human biosafety level 3 laboratories at Wageningen Bioveterinary Research, Lelystad, The Netherlands.

### Virus

SARS-CoV-2/human/NL/Lelystad/2020 was isolated from a human subject early in the pandemic and used for challenge inoculation. Details can be found in Gerhards et al. (2021) and the sequence is available under GenBank accession number MZ14458.

### Animal housing and experimental design

Eight male and eight female cats were obtained from Marshall Bioresources. They were four months of age, weighed between 1.8 and 2.8kg upon arrival, were raised as SPF cats and vaccinated against rabies and feline rhinotracheitis-calici-panleukopenia. The acclimatization time was 11 days and the acclimatization time was also used to further socialize the animals to the animal technical staff. No blinding of personnel took place, and only healthy animals were included in the study.

Using the randomizer function of Microsoft Excel, cats were first divided by gender and then attributed to one of four experimental units with four cats per group [group 1 (males), 2 (females), 3 (males) and 4 (females)]. Each group was housed in a separate hBSL3 animal unit, which in turn was divided in two pens A and B. Each pen had a volume of approximately 13.5m^3^ (1.9m width, 2.75m length and 2.58m height). Of each group (X=1, 2, 3 or 4), cats X.1 and X.2 were housed in pen A, and the other two cats in pen B (cats X.3 and X.4) until study day 6. On study day D0, cat X.2 was temporarily removed from pen A for one day, and placed back on D1. On D6, cats X.1 and X.2 were removed from pen A and placed in pen B, while cats X.3 and X.4 were removed from pen B and placed in pen A until the end of the study (D23). In every animal unit, pen A and B were separated by more than 30cm distance and a plastic separator to prevent droplet transmission between the pens. To avoid transmission by personnel, animal technicians first entered pen B before approaching pen A until D6, and from D7 onwards pen A first, and pen B second. The inoculated cats, i.e. X.1, were handled last. No exchange of equipment took place between pens. The average temperature throughout the study was 20.4°C in group 1 (range 19.9 - 21.1), 19.9°C in group 2 (range 18.3 - 21.0), 19.6°C in group 3 (range 19.0 – 20.3) and 18.7°C in group 4 (range 17.9 – 19.4). Relative humidity throughout the study was on average 51.3% in group 1 (range 43.0 – 68.0), 52.6% in group 2 (range 39.0-76.0), 52.5% in group 3 (range 42.0 - 71.0) and 63.0% in group 4 (range 46.0 – 77.0). There were 12 hours light and 12 hours dark per day. Cats were provided with water and commercial cat pellets ad libitum. Feed was removed from the pens on the evenings before the days cats underwent anaesthesia. Several types of commercial cat toys were available as enrichment, as well as hammocks, baskets, elevated resting spots, pillows, towels and scratching posts.

### Animal inoculation and samplings

All cats were anesthetized on D0, 8 and 15 by intramuscular injection of 0.04mg/kg medetomidine and 4mg/kg ketamine, which was antagonized by 0.1mg/kg atipamezole intramuscularly. Anaesthetics were injected in the hind legs, and the procedures were performed in the animal units.

On D0 in the morning, cats 1.1, 2.1, 3.1 and 4.1 were inoculated intranasally with 10^4.5^ TCID_50_ SARS-CoV-2 under general anaesthesia in a volume of 0.5mL (0.25mL per nostril, pipetted dropwise synchronous with the cat’s breathing rhythm). Every cat was inoculated by a different animal technician. Cats were observed daily for their general health and respiratory signs post challenge. Body weight was measured regularly (D-11, −10, −7, −5, −3, 0, 2, 4, 7, 9, 11, 14, 16, 18, 21 and 23).

Nasal swabs and oropharyngeal swabs, as well as 2mL serum and 1mL heparinized blood were taken under general anaesthesia on D0, 8, 15 and upon euthanasia. Oral and rectal swabs were taken without anaesthesia on D0, 2-5, 7, 9-11, 14, 16-18, 21 and upon euthanasia. Electrostatic dust cloths were used to sample the environment daily from D0 to 23. There were five different sample locations: floor (random area of 10×10cm), wall (random area of 1×1m), litter box (plastic, rim of one litter box was wiped), feeding tray (stainless steel, one feeding tray was wiped before feed was added), toy (cotton and plastic, a random toy was wiped).

### Pathology and immunohistochemistry

Cat 4.3 died on D18 unrelated to SARS-CoV-2 infection and was necropsied the same day. On D23, the remaining 15 cats were euthanized to perform a full macroscopic examination of all major organs and to collect blood and respiratory tract samples. Euthanasia was performed by intramuscular injection of anaesthetics (0.06mg/kg medetomidine and 6mg/kg ketamine) in the hind leg, followed by blood sampling through the aorta and exsanguination through the brachiocephalic vein. The lungs were weighed after exsanguination, and expressed as percentage of the body weight on the day of necropsy. Tissue samples of the lungs (inflated with formaldehyde), nasal conchae, trachea, tracheobronchial lymph node, intestine (duodenum, ileum and colon), mesenterial lymph node and pancreas were taken in 4% neutral buffered formaldehyde for histopathology/immunohistochemistry (IHC) and frozen for virological analyses (lungs and conchae).

Formalin-fixed samples were embedded in paraffin, sectioned at 5μm and stained with haematoxylin and eosin (H&E) for histological examination according to general pathology principles. IHC of histological specimens was performed with an antibody directed against the SARS-CoV nucleoprotein, as previously described (Gerhards et al. 2021).

### Organ suspensions, environmental samples and swabs

Lung and conchae samples were stored at ≤-70°C until further processing. Each sample was weighed before homogenization in 6mL of MEM, supplemented with 1% antibiotic/antimycotic solution (both from Gibco; Thermo Fischer Scientific; Waltham, MA, USA) in an Ultra Turrax Tube Drive (IKA; Staufen, Germany) for 30s (lungs) or 50s (conchae) at 6000rpm. Subsequently, homogenates were cleared by centrifugation at 3400×*g* at 4°C for 15min. Cleared suspensions were mixed 1:3 with Trizol-LS (Sigma; Saint Louis, MO, USA) and stored at ≤-15°C until RNA isolation.

Environmental tissues were collected in individual seal bags and frozen directly at −70°C. To recover viral RNA, 10mL of tissue culture medium (MEM) was added to the tissues and squidged vigorously, before taking a sample of 85μL in 255μL Trizol-LS for RNA isolation.

Following sampling, all swabs were directly submerged in approximately 2mL of tissue culture medium (MEM), kept on melting ice before vigorously vortexing for 30s on a vortex (Labdancer, VWR International B.V., Amsterdam, the Netherlands), followed by centrifugation for 5 minutes at 1500×*g* and 4°C. One sample aliquot of 200μL was mixed with 200μL lysis buffer (Molgen, Veenendaal, the Netherlands), and stored at −15°C until RNA isolation.

### RNA extraction and PCR

RNA of samples stored in Trizol-LS was extracted using Direct-zol™ RNA MiniPrep kit (Zymo Research, RefNo R1013; Irvine, CA, USA) according to manufacturer’s instructions, without DNase treatment. RNA of samples stored in Molgen lysis buffer was isolated by an automated robot system (PurePrep 96) using the Molgen RNA isolation kit (OE00290096). Isolated RNA was stored at −70°C.

Viral RNA was detected with a primer and probe set annealing to the E gene according to (Corman et al. 2020). Subgenomic viral RNA was detected with a prime and probe set as described in (Wolfel et al. 2020). PCR results were expressed as Log10 viral RNA copies, based on standard curves, as described previously (Gerhards et al. 2021).

Because of the high Ct values, all swabs were isolated and tested by E-gen PCR two times. Swab samples that yielded a Ct value only once were isolated and tested a third time. Only those swabs that tested positive twice were reported as positive samples and were subsequently tested by subgenomic PCR.

### Detection of neutralizing antibodies

Collected blood samples were separated by centrifugation for 10min at 1250×*g* at room temperature (RT) after clotting for at least 1h at RT. Resultant sera were stored at ≤-15°C before heat inactivation for 2h at 56°C.

Wildtype virus neutralization tests (VNT) and immuno-Peroxidase Monolayer Assay (IPMA) were performed on Vero E6 cells (ATCC® CRL-1586™; Manassas, VA, USA) in technical duplicates by 3-fold serial dilutions on 96-well plates, as described previously (Gerhards et al. 2021). Briefly, serum was initially diluted 1:10 and 50μL of each sample were added to 50μL SARS-CoV-2 (~100 TCID_50_) in MEM. After an incubation step of 1.5h, 15’000 Vero-E6 cells were added to each well. After 4 days, plates were fixed with 4% formaldehyde and permeabilized with ice-cold 100% methanol.

To visualize SARS-CoV-2, plates were additionally permeabilized with 1% Triton X-100 and unspecific binding was blocked with 5% normal horse serum in PBS. A rabbit antiserum (rabbit-anti-SARS-CoV-2-S1-2ST (619F), Davids Biotechnologie GmbH) was used to detect the viral spike protein. After incubation with goat-anti-rabbit-HRP (Dako; Agilent; Santa Clara, CA, USA), followed by addition of AEC (3-Amino-9-ethylcarbazole) substrate solution, a clear red-brown colour developed within 30–40 minutes, which was subsequently evaluated under a standard light microscope.

The titre of each duplicate was calculated as the average of the reciprocal value of the last dilution that showed at least 50% neutralization, after log transformation. The titers are expressed as virus microneutralization titer 50 (MN50).

### ELISA

Anti-S1 ELISA of sera was performed as described previously (Zhao et al. 2021). Briefly, sera were diluted 1:50, added to S1-protein coated microtiter plates and incubated for 1h at 37°C before incubation with horseradish-peroxidase conjugated secondary goat-anti-cat IgG/HRP (dilution 1:4000, Rockland Immunochemicals Inc., Limerick, PA, USA) for 1h at 37°C. The reaction was visualized using 3,3’,5,5’-tetramethylbenzidine, quenched by sulfuric acid and optical density measurement at 450nm.

### PBMC isolation and interferon-γ-ELISpot

To isolate peripheral blood mononuclear cells (PBMCs), 15mL ficoll (GE Healthcare Bio Sciences B.V. Sweden) was added to leucosep tubes (GREINER BIO-ONE B.V., Germany), and spun down for 1 minute at 1.000x*g*. Fifteen mL of heparinized blood was diluted 1:1 with RPMI1640 (Life Technologies, The Netherlands) and added to the leucosep tubes containing ficoll. After centrifugation for 10 minutes at 1000x*g*, the interface was harvested, diluted 1:1 with cell culture medium and spun down for 7 minutes at 400x*g*. Next, cells were incubated with 4500μL ACK lysis buffer (Life Technologies, USA) for 5 minutes at room temperature and spun down again for 7 minutes at 400x*g*. The resultant pellet was resuspended in 10mL cell culture medium and cells were counted. 2×10^5^ cells were added to an ELISpot plate, prepared according to manufacturer’s instructions (Cat IFN-γ ELISpot Plus Kit, MABTech, Nacka Strand, Sweden) containing 100uL culture medium with or without stimulus. The following stimuli were used: Spike protein 30μg/mL (SARS-CoV-2 (2019-nCoV) Spike Protein (S1+S2 ECD, His tag), Sino Biological, Eschborn, Germany), nucleoprotein 10μg/mL (SARS-CoV-2 (2019-nCoV) NucleocapsidProtein (His tag), Sino Biological, Eschborn, Germany), SARS-CoV-2 Spike protein peptide pool 6μg/mL (Pepscan Presto, Lelystad, the Netherlands), concanavalin A 20μg/mL (Merck, Germany). The cells were then incubated for 48 hours at 37°C + 5%CO2 and spots were developed according to manufacturer’s instructions. Spots were analysed by a ELISpot reader (VSpot Spectrum, AID GmbH, Germany).

### Statistical analyses incorporating environmental transmission

The observed number of cases in an interval in each separated pen is the dependent variable. This dependent variable is assumed to be binomial distributed with the number of non-infected cats as the binomial total. Additionally, it is assumed that inoculated and contact infected cats are equally infectious. In this case, analysis can be done as Bernoulli trials for each recipient separately. Whether the recipient becomes infected also depends on the number of infected cats in the pen through the infectivity present in the environment. The environmental load is assumed to change deterministically according to the following differential equation: 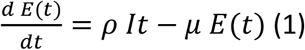

Note that “It” is always a whole number and changes stochastically with integer jumps. Hence the different notation than for E(t) which changes continuously. Given the “It” and “E0” (the environmental contamination at the start of the interval) values E(t) can be calculated during each interval by solving equation (1). Without loss of generality the value of *ρ* can be fixed. Here we will do this in such a way that 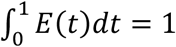 if the environment is not yet contaminated at the start of the interval. For an interval Δt the probability to become infected for each of the recipient cats is: 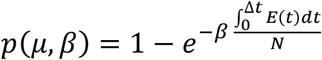. The parameters *μ* and *β* can be estimated by maximum likelihood either directly by optimizing the log likelihood as we did here or by Generalised Linear Models using an offset (Velthuis et al. 2002). More details of the formulas used are given in Supplementary material 3.

## Acknowledgements

This study was commissioned by the Dutch Ministry of Agriculture and Food under the ARVODI-2018 agreement concerning policy supporting research on Covid-19 in cats and dogs. We are thankful to Rineke de Jong, Stéphanie Vastenhouw, Sophie van Oort, José Harders-Westerveen, Bregtje Smid, Rianka Vloet, Romy Dresken, Julia Antonissen, Judith Bonsing, Ralph Kok, Pieter Roskam, Patrik Brandenburg and Corry Dolstra for their excellent technical contribution. We thank the staff of WBVR’s animal facility and biosafety officers.

We furthermore thank Berend Jan Bosch and Wentao Li (University of Utrecht, The Netherlands) for providing us the anti-spike S1 antibody used in this study for staining of the cell monolayers in the IPMA.

Illustrations in Figure 1 were created with BioRender.com.

