## Supplemental material 1 and 2 for "Efficient direct and limited environmental transmission of SARS-CoV-2 lineage B.1.22 in domestic cats"

*[a]Department of Bioinformatics, Epidemiology and Animal models, Wageningen Bioveterinary Research, Lelystad, The Netherlands; [b]Department of Bacteriology, Host-Pathogen Interactions and Diagnostic Development, Wageningen Bioveterinary Research, Lelystad, The Netherlands; [c]Division of Pathology, Faculty of Veterinary Medicine, Utrecht University, The Netherlands; [d]Department of Animal Breeding and Genomics, Wageningen Research, The Netherlands; [e]Division Infectious Diseases and Immunology, Section Virology, Veterinary Faculty, Utrecht University, The Netherlands; [f]Department of Population Health Sciences, Veterinary Epidemiology, Veterinary Faculty, Utrecht University, The Netherlands; [g]Department of Virology and Molecular Biology, Wageningen Bioveterinary Research, Lelystad, The Netherlands [h]Department of Quantitative Veterinary Epidemiology, Wageningen University, The Netherlands*

#### Supplemental material

#### Supplemental material S1: Activity analysis

To investigate the effect of SARS-CoV-2 on activity, video analysis was performed. A RGB camera was placed in pens A of groups 1, 2, 3 and 4 to record daily sessions of 19 hours, from 3 p.m. to 10 a.m. the next day (1920 x 1080 pixels, 30 fps). Recordings of DPI -6 until DPI 5 were analyzed, of which 5 recordings were discarded (3 corrupted recordings, one video on which a cat had escaped from the pen (before inoculation) and one recording with a flickering light leading to disturbance). Of each recording, the first 2.5 hours and the last 0.5 hour were discarded, because animal caretakers entered the pens during these times. In addition, 2 minutes were discarded around the timestamps that the camera changed from day to night vision (~5 hours into the video) and from night to day vision (~17 hours into the recording).

Activity was quantified using pixel differences. A sampling rate of 1 out of 5 frames was applied, so that 6 frames per second were kept for the analysis. Frames were converted to grayscale and for each pair of subsequent frames, intensity values (ranging from 0 to 255) were compared. A pixel was considered to change when its intensity value differed by more than 40 between subsequent frames (see Figure S1 for an example). To remove extreme outliers caused by e.g. light flashes, a moving median of 6 frames was used. Results of the 6 days before inoculation were then compared to those in the 5 days after inoculation. Analyses were performed in Python with the OpenCV set of open source computer vision tools (Bradski, 2000).

Activity results showed a circadian rhythm (Figures S2-S5). Activity was relatively low during the night and peaked around dusk (3-6 hours into each video) and dawn (15-18 hours into each video). This is in line with the crepuscular nature of domestic cats (Cove et al., 2018; Parker et al., 2019).

There was no difference in activity at pen level between the 6 days before inoculation and the 5 days after inoculation (Figure S6). When comparing the 6 days before inoculation with 2 days after inoculation, instead of 5 days after inoculation, there was also no observable difference. The absence of change in activity post inoculation is in line with the absence of clinical symptoms in this study.

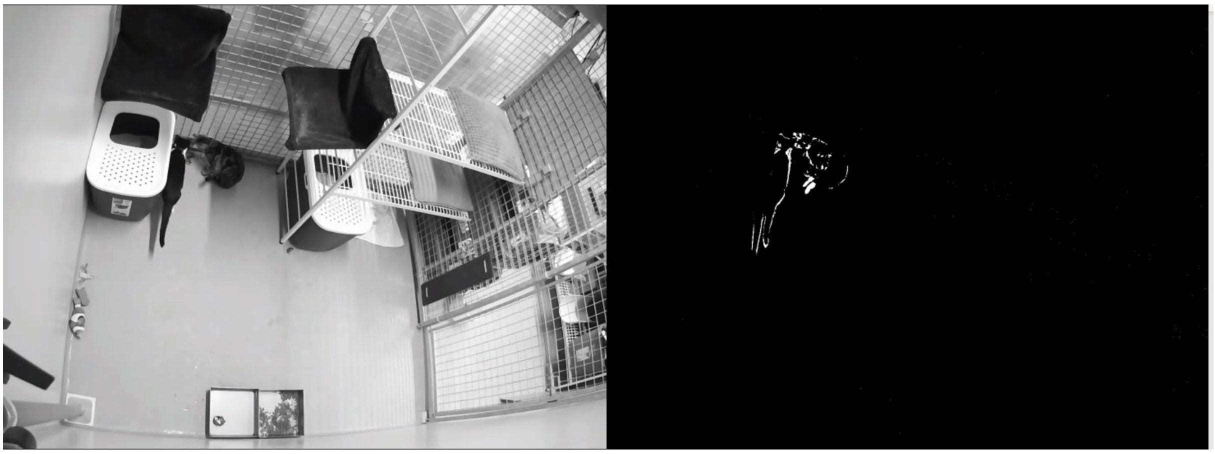

**Figure S1.** Quantifying activity with pixel differences. Left: grayscale frame of original video. Right: pixels that changed more than 40 in intensity values between current frame and previous frame shown in white (rest is black).

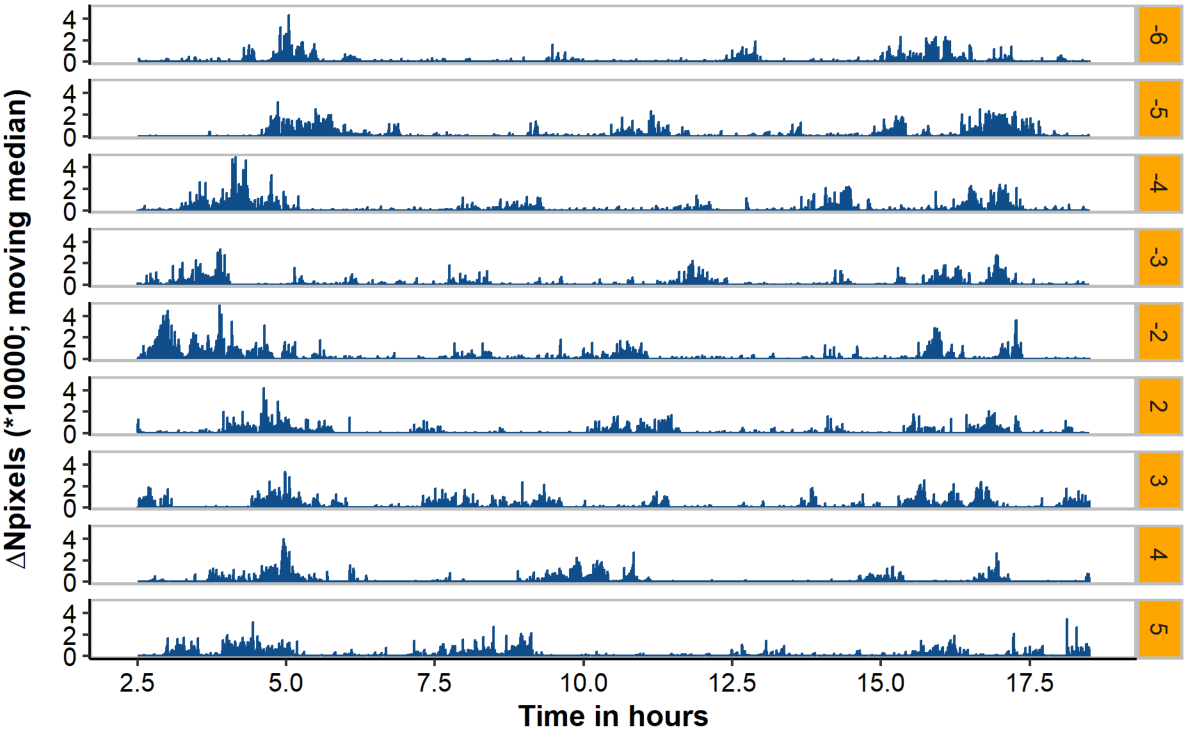


**Figure S2.** Moving median of the number of pixels that differed between subsequent frames (ΔNpixels) for the days before and after inoculation for pen A. Note: the videos for day -1 and day 1 were unavailable/discarded.

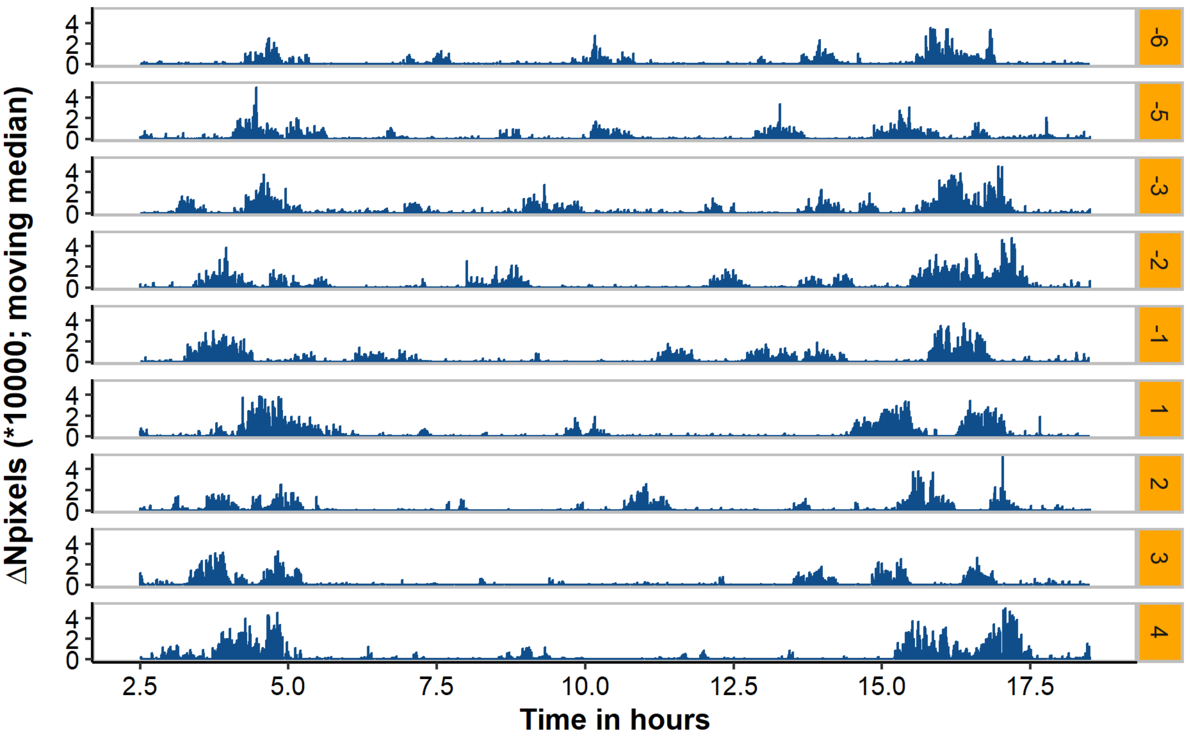


**Figure S3.** Moving median of the number of pixels that differed between subsequent frames (ΔNpixels) for the days before and after inoculation for pen B. Note: the videos for day -4 and day 5 were unavailable/discarded.

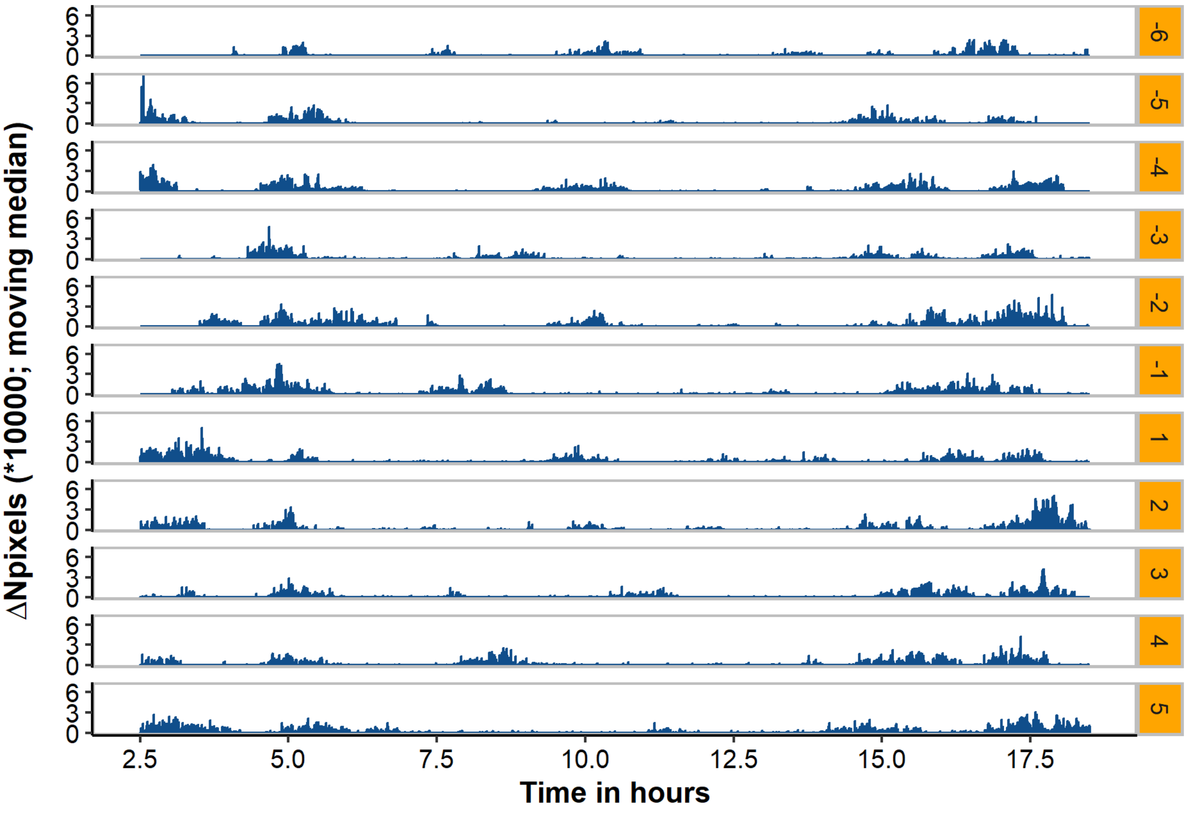


**Figure S4.** Moving median of the number of pixels that differed between subsequent frames (ΔNpixels) for the days before and after inoculation for pen C.

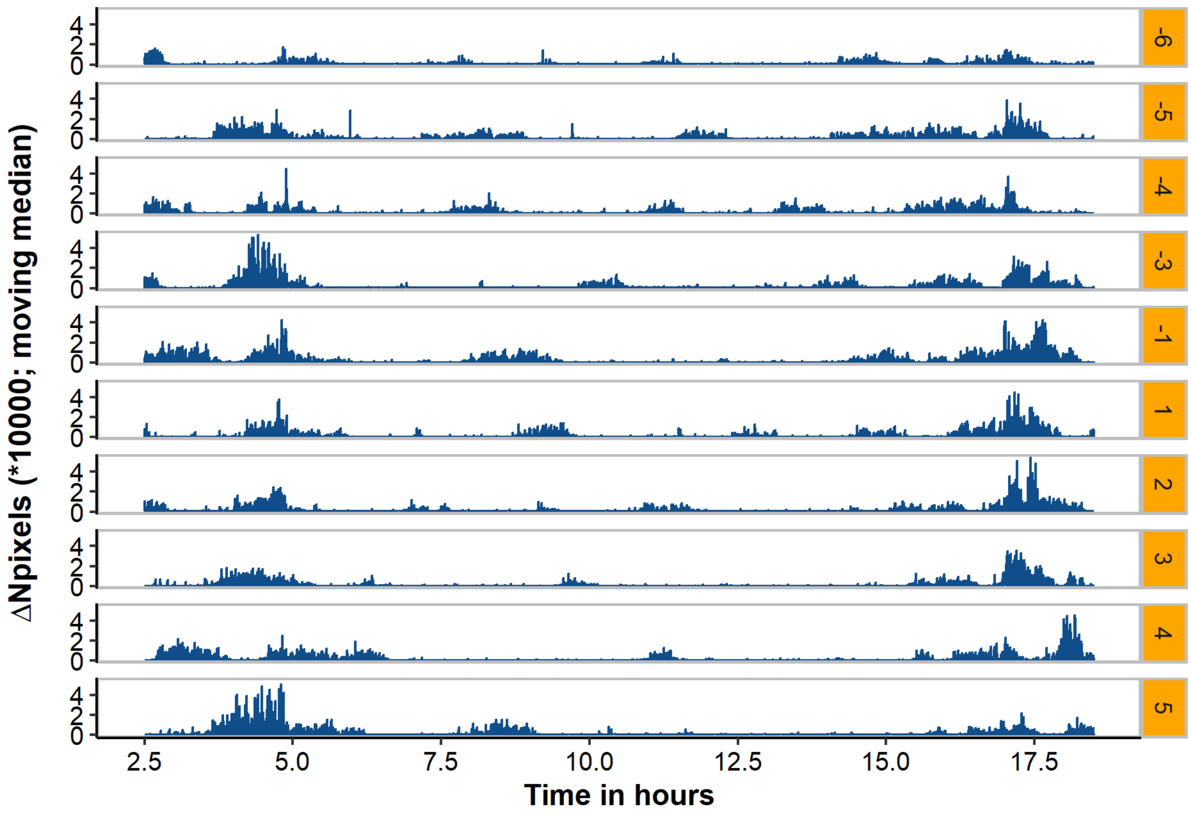


**Figure S5.** Moving median of the number of pixels that differed between subsequent frames (ΔNpixels) for the days before and after inoculation for pen D. Note: the video for day -2 was unavailable.

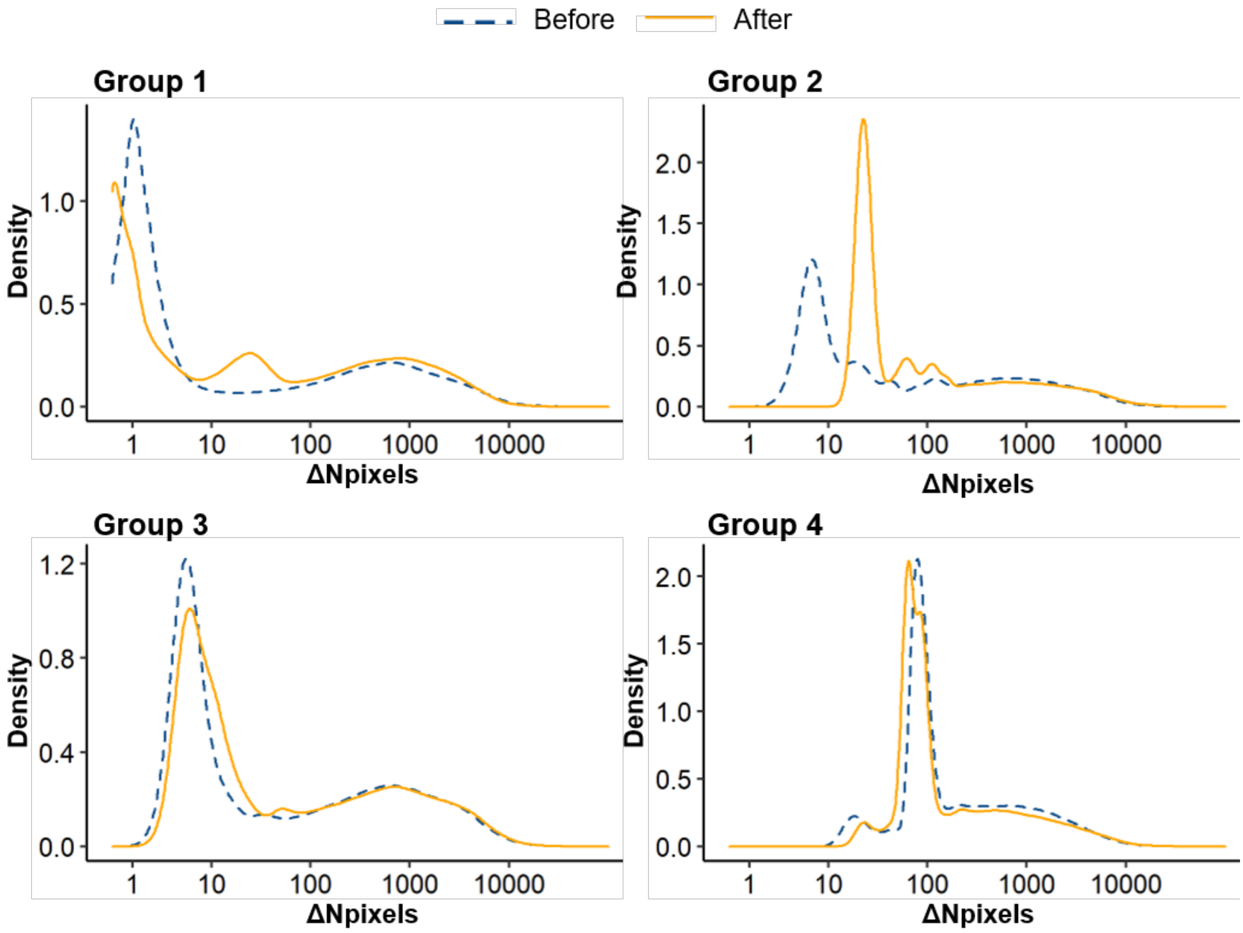


**Figure S6.** Density plots of the number of pixels that differed between subsequent frames (ΔNpixels) for the 6 days before inoculation and 5 days after inoculation for groups 1, 2, 3 and 4. Note: a (pseudo) $^{10}$log-scale was used for the x-axis.

#### Supplemental material S2: Mathematical analysis

**Datasets**

Data 1

E-gene PCR as indication of infectivity for each infected animal. Infection determined by seroconversion. This is analyzing only the first part of the experiment (direct contact).

| "cat ID" | "start" | "end" | "#S" | "prev" | "case" | "history" |
|---|---|---|---|---|---|---|
| 1.2 | 1 | 5 | 1 | 0.5 | 0 | { } |
| 1.2 | 5 | 6 | 1 | 0.5 | 1 | {{0.5,1,5}} |
| 2.2 | 1 | 5 | 1 | 0.5 | 0 | { } |
| 2.2 | 5 | 6 | 1 | 0.5 | 1 | {{0.5,1,5}} |
| 3.2 | 1 | 6 | 1 | 0.5 | 0 | { } |
| 4.2 | 1 | 6 | 1 | 0.5 | 0 | { } |

Data 2

E-gene PCR as indication of infectivity for each infected animal. Infection determined by seroconversion.

| "cat ID" | "start" | "end" | "#S" | "prev" | "case" | "history" |
|---|---|---|---|---|---|---|
| 1.2 | 1 | 5 | 1 | 0.5 | 0 | { } |
| 1.2 | 5 | 6 | 1 | 0.5 | 1 | {{0.5,1,5}} |
| 2.2 | 1 | 5 | 1 | 0.5 | 0 | { } |
| 2.2 | 5 | 6 | 1 | 0.5 | 1 | {{0.5,1,5}} |
| 1.3 | 6 | 13 | 1 | 0 | 0 | {{0.5,1,6}} |
| 1.4 | 6 | 23 | 1 | 0 | 0 | {{0.5,1,6}} |
| 2.3 | 6 | 23 | 1 | 0 | 0 | {{0.5,1,6}} |
| 2.4 | 6 | 23 | 1 | 0 | 0 | {{0.5,1,6}} |
| 3.3 | 6 | 23 | 1 | 0 | 0 | {{0.5,1,6}} |
| 3.4 | 6 | 23 | 1 | 0 | 0 | {{0.5,1,6}} |
| 4.3 | 6 | 23 | 1 | 0 | 0 | {{0.5,1,6}} |
| 4.4 | 6 | 23 | 1 | 0 | 0 | {{0.5,1,6}} |
| 3.2 | 1 | 6 | 1 | 0.5 | 0 | { } |
| 4.2 | 1 | 6 | 1 | 0.5 | 0 | { } |

Data 3

E-gene PCR as indication of infectivity for each infected animal. Infection determined by SG PCR positivity

| "cat ID" | "start" | "end" | "#S" | "prev" | "case" | "history" |
|---|---|---|---|---|---|---|
| 1.2 | 1 | 5 | 1 | 0.5 | 0 | { } |
| 1.2 | 5 | 6 | 1 | 0.5 | 1 | {{0.5,1,5}} |
| 2.2 | 1 | 5 | 1 | 0.5 | 0 | { } |
| 2.2 | 5 | 6 | 1 | 0.5 | 1 | {{0.5,1,5}} |
| 4.2 | 1 | 2 | 1 | 0.5 | 1 | { } |
| 1.3 | 6 | 7 | 1 | 0 | 1 | {{0.5,1,6}} |
| 1.4 | 6 | 7 | 1 | 0 | 0 | {{0.5,1,6}} |
| 1.4 | 7 | 13 | 1 | 0.5 | 0 | {{0.5,1,6}} |
| 1.4 | 13 | 23 | 1 | 0 | 0 | {{0.5,1,6},{0.5,7,13}} |
| 2.3 | 6 | 23 | 1 | 0 | 0 | {{0.5,1,6}} |
| 2.4 | 6 | 23 | 1 | 0 | 0 | {{0.5,1,6}} |
| 3.3 | 6 | 23 | 1 | 0 | 0 | {{0.5,1,6}} |
| 3.4 | 6 | 23 | 1 | 0 | 0 | {{0.5,1,6}} |
| 4.3 | 6 | 23 | 1 | 0 | 0 | {{0.5,1,6},{0.5,2,6}} |
| 4.4 | 6 | 23 | 1 | 0 | 0 | {{0.5,1,6},{0.5,2,6}} |
| 3.2 | 1 | 6 | 1 | 0.5 | 0 | { } |

Data4

E-gene PCR positivity as indication of infectivity and as indication of infection.

| "cat ID" | "start" | "end" | "#S" | "prev" | "case" | "history" |
|---|---|---|---|---|---|---|
| 1.2 | 1 | 5 | 1 | 0.5 | 0 | { } |
| 1.2 | 5 | 6 | 1 | 0.5 | 1 | {{0.5,1,5}} |
| 2.2 | 1 | 5 | 1 | 0.5 | 0 | { } |
| 2.2 | 5 | 6 | 1 | 0.5 | 1 | {{0.5,1,5}} |
| 3.2 | 1 | 2 | 1 | 0.5 | 1 | { } |
| 4.2 | 1 | 2 | 1 | 0.5 | 1 | { } |
| 1.3 | 6 | 7 | 1 | 0 | 1 | {{0.5,1,6}} |
| 1.4 | 6 | 7 | 1 | 0 | 0 | {{0.5,1,6}, {0.5,7,8}} |
| 1.4 | 7 | 8 | 1 | 0.5 | 0 | {{0.5,1,6}, {0.5,7,8}} |
| 1.4 | 8 | 9 | 1 | 0.5 | 1 | {{0.5,1,6}, {0.5,7,8}} |
| 3.4 | 6 | 8 | 1 | 0 | 0 | {{0.5,1,6}, {0.5,2,6}} |
| 3.4 | 8 | 9 | 1 | 0 | 1 | {{0.5,1,6}, {0.5,2,6}} |
| 4.3 | 6 | 7 | 1 | 0 | 1 | {{0.5,1,6}, {0.5,2,6}} |
| 4.4 | 6 | 7 | 1 | 0 | 1 | {{0.5,1,6}, {0.5,2,6}} |
| 2.3 | 6 | 23 | 1 | 0 | 0 | {{0.5,1,6}} |
| 2.4 | 6 | 23 | 1 | 0 | 0 | {{0.5,1,6}} |
| 3.3 | 6 | 8 | 1 | 0 | 0 | {{0.5,1,6}, {0.5,2,6}} |
| 3.3 | 8 | 9 | 1 | 0.5 | 0 | {{0.5,1,6}, {0.5,2,6}} |
| 3.3 | 9 | 13 | 1 | 0 | 0 | {{0.5,1,6}, {0.5,2,6}, {0.5,8,9}} |

**Algorithm**

### Definitions

```
Clear[HazardEt, IntegratEt, E0, R0, totalE, Exposure,
 datalinelikelihood, logdatalikelihood, profilelikelihoodβ,
 profilelikelihoodμ, profilelikelihoodR, profileβ, profileμ, profileR]
```

```
(* environmental contamination D[Env[t],t]==
 ϕ(μ) It - μ E(t) with ϕ(μ)standardised *)
```

In[24]:=
```
(* instantaneous hazard rate when It is constant
 in the interval and the amount is E0 at the start*)
```

In[25]:=

$$\texttt{HazardEt[t\_, It\_, E0\_, }\mu\texttt{\_] :=} \frac{\mu\left(1 - e^{-\mu t}\right)}{-1 + e^{-\mu} + \mu} \texttt{It} + e^{-\mu t}\,\texttt{E0}$$

```
(* accumulated over time period when It is constant during
  the interval and E0 is the amount at the start of the interval
  Note: the standardisation makes it equal to It for
        t=1 and E0=0 *)
```

In[27]:=

$$\texttt{IntegratEt[t\_, It\_, E0\_, }\mu\texttt{\_] :=} \frac{-1 + e^{-\mu t} + \mu t}{-1 + e^{-\mu} + \mu} \texttt{It} + \frac{1 - e^{-\mu t}}{\mu} \texttt{E0}$$

```
(* amount already present at the start of this interval
 given historical exposure(s) with shedding from infected (I/N),
value in history[[x,1]], during previous intervals ending with history[[x,3]]
 and starting at history[[x,2]] time units and decay then continues
 until the start of the interval under consideration at t0. *)
```

In[29]:=

$$\texttt{E0[history\_, }\mu\texttt{\_, t0\_] := If}\Big[\texttt{Length[history]} \le 0, 0, \sum_{x=1}^{\texttt{Length[history]}} \big(\texttt{If[t0} \ge \texttt{history[[x, 3]], HazardEt[history[[x, 3]] - history[[x, 2]], history[[x, 1]], 0, }\mu\texttt{]} * \texttt{Exp[-}\mu\texttt{ (t0 - history[[x, 3]])], 0]}\big)\Big]$$

In[ ]:=

$$\texttt{totalE[}\mu\texttt{\_, It\_] :=} \frac{\mu}{-1 + e^{-\mu} + \mu} \texttt{It}$$

In[ ]:=
```
R0[β_, μ_, T_] := β totalE[μ, 1] T
```

In[ ]:=
```
Exposure[n_, μ_, data_] :=
 IntegratEt[data[[n, 3]] - data[[n, 2]], data[[n, 5]], E0[data[[n , 7]], μ, data[[n, 2]]], μ]
```

In[ ]:=
```
(* dataline likelihood with I/N data[[n,6]] during the interval
 (length data[[n,7]])present resulting in data[[n,5]] positive individuals
 and data[[n,4]] infection negative individuals. Exposure history is
 in data[[n,8]] with three values per entry (see explanantion of E0) *)
```

In[ ]:=

$$\texttt{datalinelikelihood[n\_, }\beta\texttt{\_, }\mu\texttt{\_, data\_] := If}\Big[\texttt{Exposure[n, }\mu\texttt{, data]} > \texttt{10\^{}-10, }\Big(\big(\texttt{1 - Exp[-}\beta\texttt{ Exposure[n, }\mu\texttt{, data]]}\big)^{\texttt{data[[n,6]]}}\ \texttt{Exp[-}\beta\texttt{ Exposure[n, }\mu\texttt{, data]]}^{\texttt{If[data[[n,6]]>0,0,1]}}\Big)\texttt{, (1; If[data[[n, 6]] > 0, Print[" Dataline ", n, " not possible"]])}\Big]$$

```
In[-]:= (* for likelihood optimisation we add the
     Log of all the observed independent likelihoods *)

In[-]:= logdatalikelihood[β_, μ_, data_] := Sum[Log[datalinelikelihood[n, β, μ, data]], {n, 1, Length[data]}]

In[-]:= (* for profile likelihood we calculate the
     parameter β given the value of μ or vice versa *)

In[-]:= profileμ[μ_, data_] :=
     β /. Flatten[FindMaximum[{logdatalikelihood[β, μ, data], β > 0}, {β, 0.1}]][[2]]

In[-]:= profileβ[β_, data_] :=
     μ /. Flatten[FindMaximum[{logdatalikelihood[β, μ, data], μ > 0}, {μ, 0.1}]][[2]]

In[-]:= profileR[R_, T_, data_] := {R / (totalE[μ, 1] T), μ} /. Flatten[
       FindMaximum[{logdatalikelihood[R / (totalE[μ, 1] T), μ, data], μ > 0}, {μ, 0.1}]][[2]]

In[-]:= (* these are then the profilelikelihoods *)

In[-]:= profilelikelihoodμ[μ_, data_] := logdatalikelihood[profileμ[μ, data], μ, data]

In[-]:= profilelikelihoodβ[β_, data_] := logdatalikelihood[β, profileβ[β, data], data];

In[-]:= profilelikelihoodR[R_, T_, data_] :=
      logdatalikelihood[profileR[R, T, data][[1]], profileR[R, T, data][[2]], data];
```

The sum in logdatalikelihood is printed as $\sum_{n=1}^{\text{Length[data]}}$.

**Profile likelihoods final model**

The profile likelihood of the decay rate parameter μ, estimated value 2.73 day$^{-1}$ (0.77-15.82)

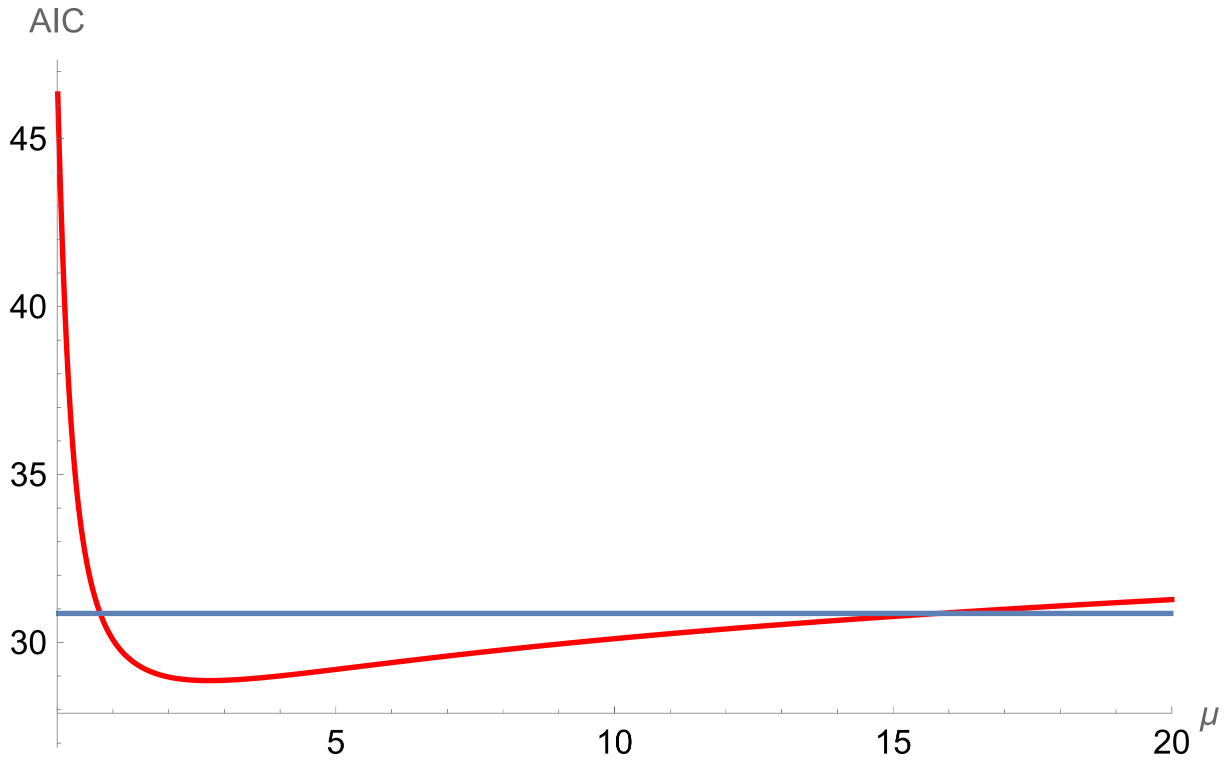


The profile likelihood of the transmission rate parameter β, estimated value 0.23 day$^{-1}$ (0.06,0.54)

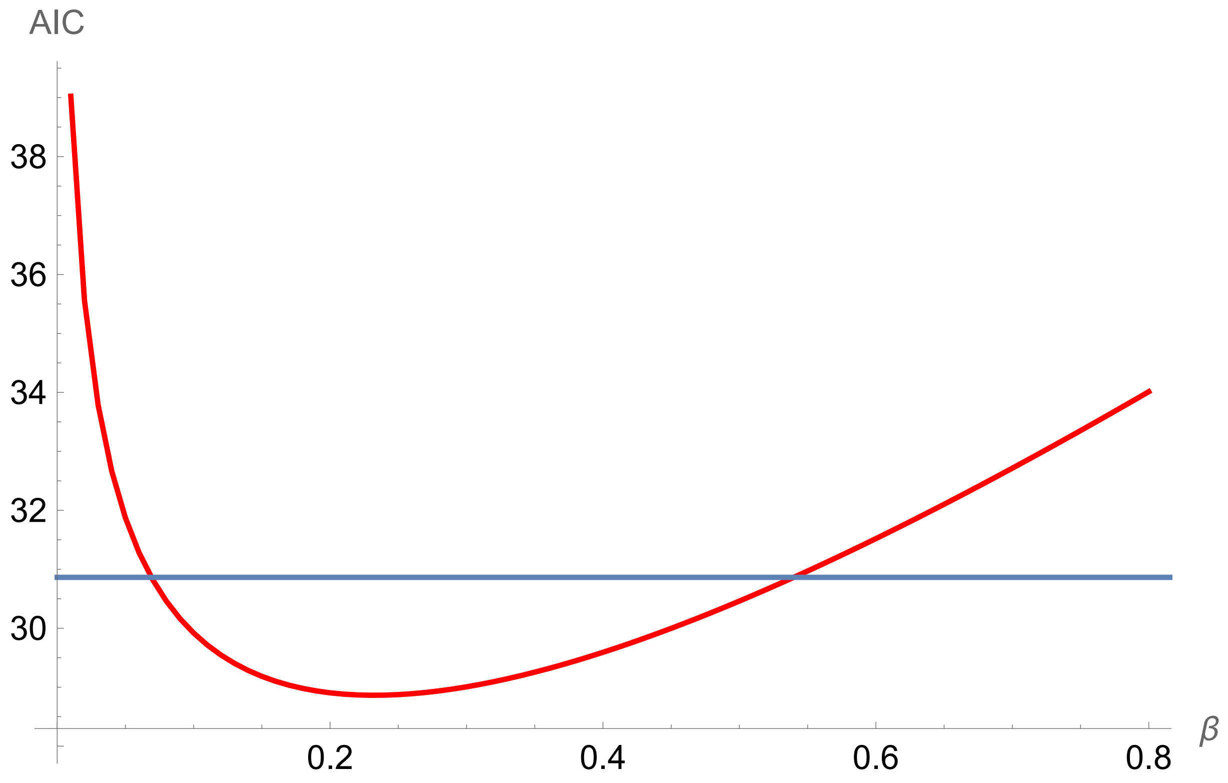

The profile likelihood of the reproduction ratio R, estimated value 2.12 (0.92-4.08).

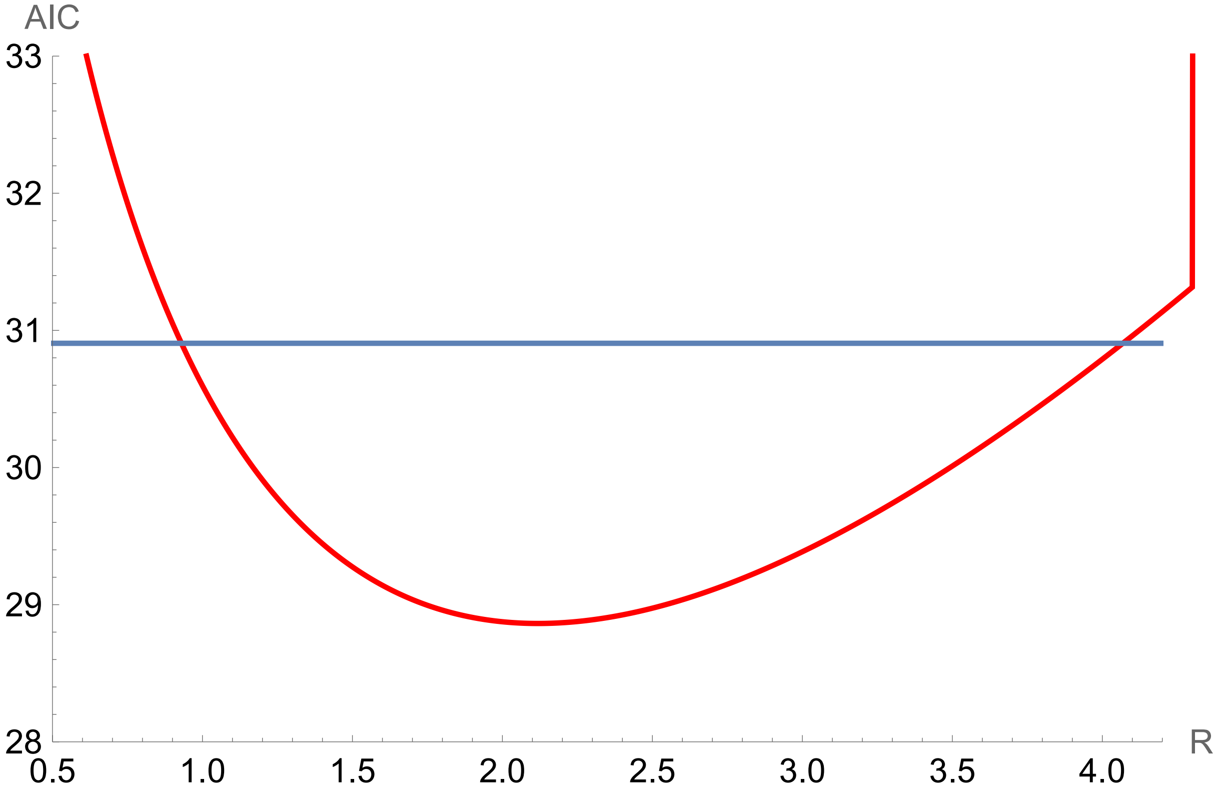
